# An *in vivo* platform for rebuilding functional neocortical tissue

**DOI:** 10.1101/2022.12.09.519776

**Authors:** Alexandra Quezada, Claire Ward, Edward R. Bader, Pavlo Zolotavin, Esra Altun, Sarah Hong, Nathan Killian, Chong Xie, Renata Batista-Brito, Jean M. Hébert

**Author notes:** Co-first authors (Alexandra Quezada, Claire Ward). Co-senior authors (Renata Batista-Brito, Jean M. Hébert).

## Abstract

Recent progress in cortical stem cell transplantation has demonstrated its potential to repair the brain. However, current transplant models have yet to demonstrate that the circuitry of transplant-derived neurons can encode useful function to the host. This is likely due to missing cell types within the grafts, abnormal proportions of cell types, abnormal cytoarchitecture, and inefficient vascularization. Here, we devised a transplant platform for testing neocortical tissue prototypes. Dissociated mouse embryonic telencephalic cells in a liquid scaffold were transplanted into aspiration-lesioned adult mouse cortices. The donor neuronal precursors differentiated into upper and deep layer neurons that exhibited synaptic puncta, projected outside of the graft to appropriate brain areas, became electrophysiologically active within one month post-transplant, and responded to visual stimuli. Interneurons and oligodendrocytes were present at normal densities in grafts. Grafts became fully vascularized by 1-week post-transplant and vessels in grafts were perfused with blood. With this paradigm, we could also organize cells into layers. Overall, we have provided proof of concept for an *in vivo* platform that can be used for developing and testing neocortical-like tissue prototypes.

## Introduction

The transplantation of pluripotent stem cell-derived neural precursors into the cortex is an exciting potential approach to repair the brain. To achieve this goal, grafted cells must re-establish damaged neural circuits that participate in the restoration of lost behavioral function. Significant progress has been made in demonstrating the feasibility of transplanting precursor cells to replace neurons in the cortex. Graft-derived neurons can survive for years in mice (Linaro et al., 2019), differentiate into appropriate neuronal subtypes that exhibit normal electrophysiological activity, project long distances outside of the graft to appropriate targets, synaptically integrate with surrounding host neurons, and respond to sensory input and participate in motor output (Espuny-Camacho et al., 2018; Falkner et al., 2016; Mansour et al., 2018; Michelsen et al., 2015; Palma-Tortosa et al., 2020; Tornero et al., 2017).

Despite these significant discoveries, it is unclear whether grafted neurons in the neocortex can encode useful behavior as a result of their electrophysiological activity. Reported behavioral benefits are instead a result of activity-independent functions such as the secretion of anti-inflammatory or neurotrophic factors (Henriques et al., 2019; Lee et al., 2007). This is unsurprising considering there are cortical cell types that are thus far missing in grafts, in addition to these grafts lacking normal cortical cytoarchitecture. While cerebral organoids display a subset of similar characteristics to a normal fetal cortex, their differentiation has thus far been abnormal after transplantation (Pham et al., 2018; Qian et al., 2020). Therefore, there is currently no method of generating facsimiles of neocortical tissue in adults, whether for the purpose of study or therapy.

Here, to address this shortcoming, we tested whether grafting cells in a three-dimensional scaffold could sustain the differentiation of all the major cortical cell types, vascularization, and a layered cytoarchitecture. Using dissociated mouse cortical fetal cells mixed with a commercial scaffold, we found that the neuronal, glial, and vascular components within the graft survived and successfully integrated with the host tissue. Our results suggest that this platform is suitable for future optimization and testing of structured, vascularized, multi-cell type neocortical tissue prototypes.

## Results

### Graft integrity is dependent on scaffold dilution

To generate consistently sized lesions, cortical tissue was removed with a biopsy punch followed by aspiration to create a cylindrical cavity that reached the corpus callosum. Biopsy punches of 1.00, 1.25, and 2.0 mm in diameter resulted in reproducible lesions (Supplementary Fig. 1A-C). For consistency, we used 1.25 mm lesions for this study unless otherwise indicated.

**Supplementary Figure 1.**
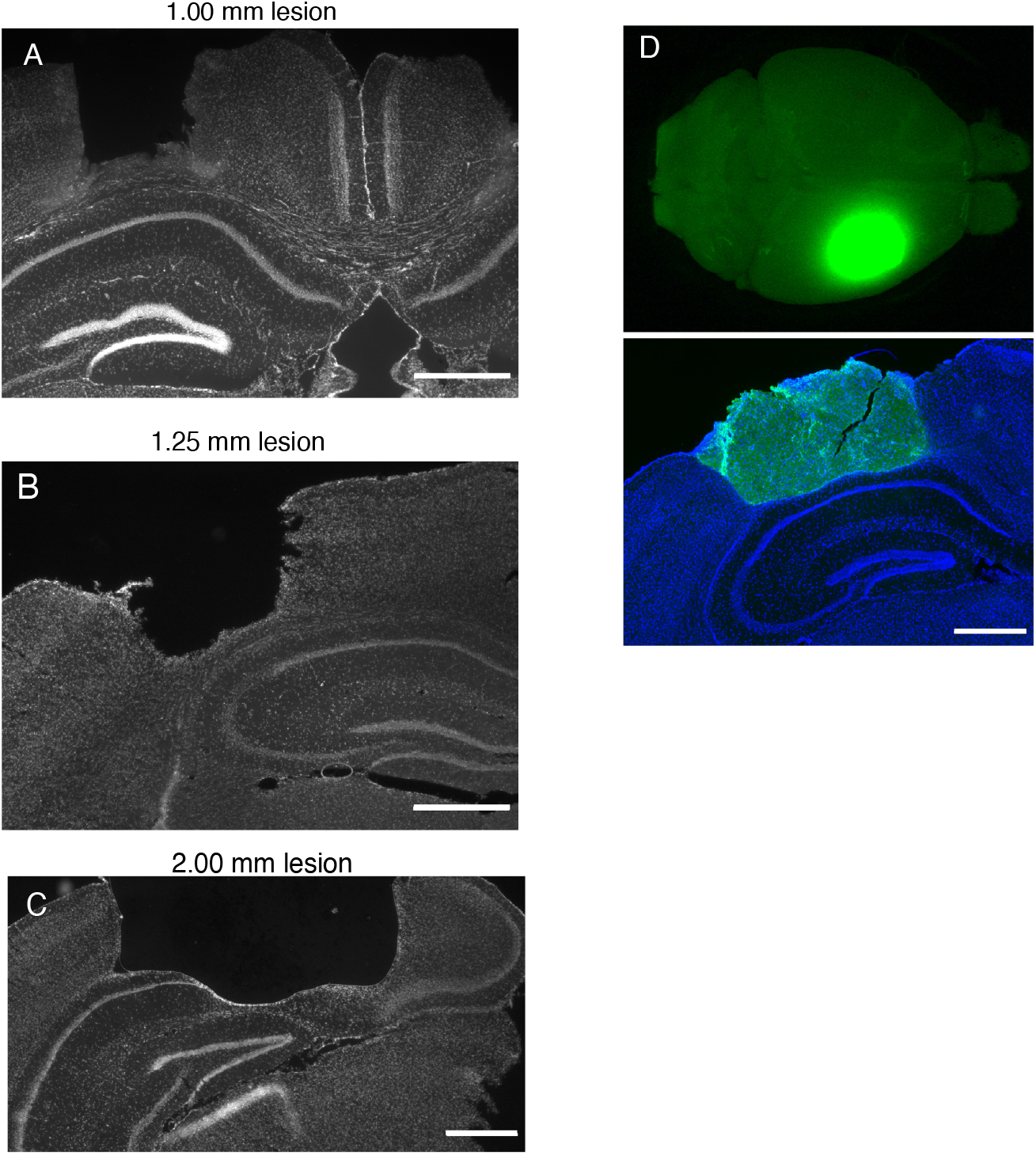
Cortical lesions can be of differing diameters. A. 1 mm cortical lesion (Scale bar=1 mm). B. 1.25 mm cortical lesion (Scale bar=1 mm). C. 2 mm cortical lesion (Scale bar=1 mm). D. Representative image of a 2mm graft. Top panel, GFP-labeled graft in whole mouse brain. Bottom panel, immunofluorescence image of coronal section of graft (Scale bar=1 mm).

To deposit and maintain cells within the lesion sites, Matrigel was chosen as the scaffold due to its suitability for neuronal and vascular differentiation and survival (Daviaud et al., 2018; Wang et al., 2020; Yang et al., 2020). To determine the optimal Matrigel dilution for transplanting mouse fetal cells, we compared Matrigel:medium dilutions of 1:2, 1:3, 1:4, and 1:5 (Supplementary Fig. 2A-B). Dilutions of 1:2 and 1:3 led to inconsistently sized grafts, often with gaps between the scaffold and the host tissue, while 1:4 diluted grafts were more consistent in size and appeared to have a more uniform donor cell distribution. The area and perimeter of grafts at 1:4 dilution (N = 7) were significantly larger than grafts with 1:3 dilution (N = 6, p =0.03 for area, 0.03 for perimeter) and 1:5 dilutions (N = 6, p = 0.04 for area, 0.02 for perimeter) (Supplementary Fig. 2C-D). Although this study was performed with 1.25 mm diameter lesions, transplants into 2 mm lesions showed that the 1:4 dilution also evenly fills larger cylindrical lesions with cells that are uniformly distributed (Supplementary Fig.1D).

The density of blood vessels (labeled as CD31+ /CD105+) within the grafts was also affected by the dilution of Matrigel (Supplementary Fig.2B, E). Grafts with a dilution of 1:3 (N = 3, p = 0.02) and 1:4 (N = 3, p = 0.002) had lower vessel density while grafts with a 1:5 dilution (N = 4, p = 0.13) were not statistically different to the density in intact contralateral cortex. This was not surprising because lower concentrations of Matrigel result in decreased matrix stiffness, allowing for more 3D structures such as vessels to form. Given the overall performance of the 1:4 Matrigel:media dilution, the remaining grafts were performed using this dilution.

**Supplementary Figure 2.**
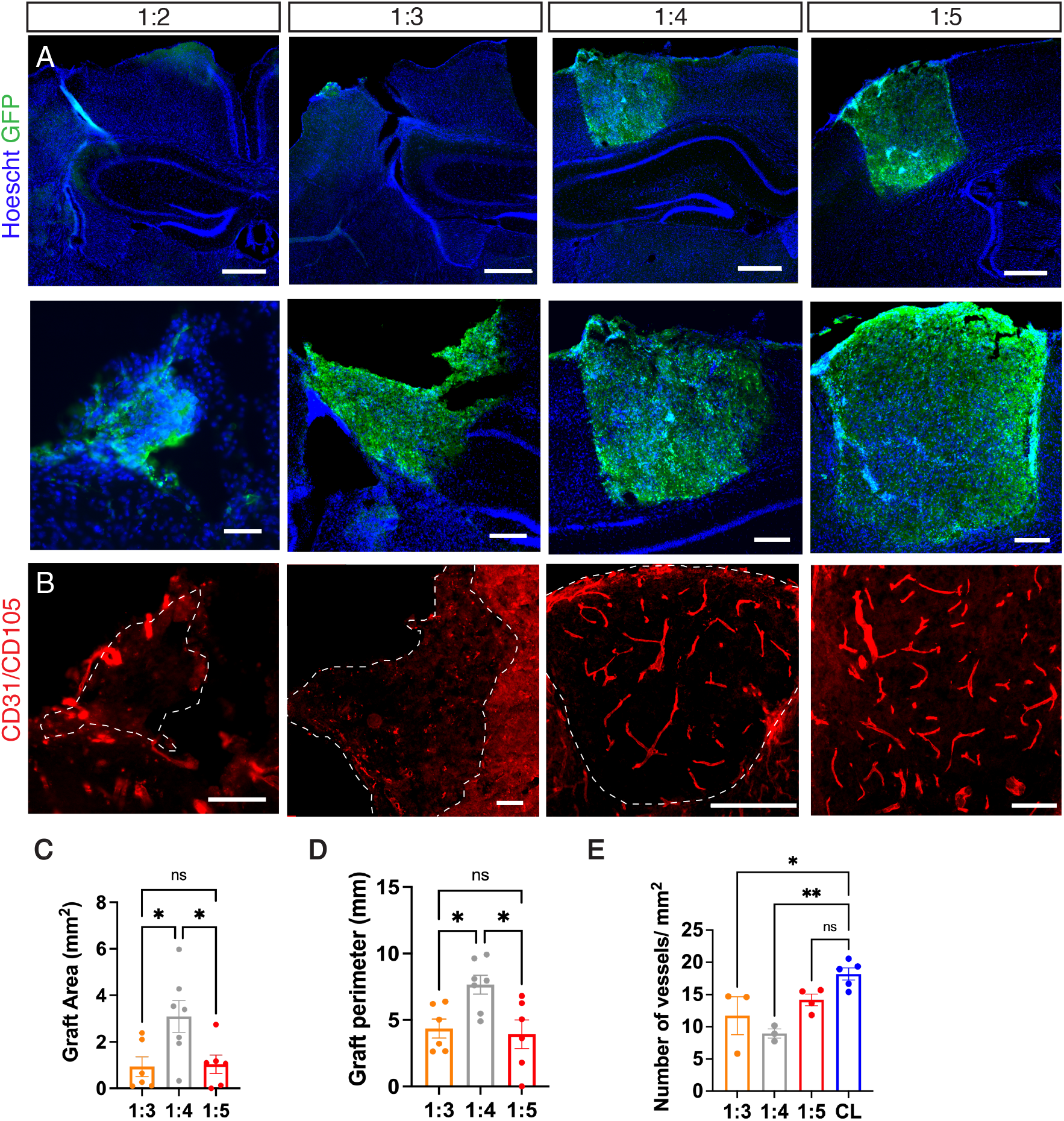
The quality of grafts is dependent on the concentration of scaffold. A. Representative immunofluorescence images of grafts with volume ratios of Matrigel:media from left to right of 1:2, 1:3, 1:4, 1:5. Top row, low magnification (Scale bars from left to right = 800 μM, 500 μM, 1 mm, 1 mm). Bottom row, higher magnification (Scale bars from left to right = 100 μM, 150 μM, 300 μM, 300 μM). B. Representative immunofluorescence images of graft vasculature for each Matrigel:media ratio (Scale bars from left to right = 100 μM, 150 μM, 500 μM, 200 μM). C. Quantification of graft area. Grafts with 1:4 dilution (N=7, mean=3.01 mm^2^), were significantly larger than 1:3, (N = 6, mean=0.93 mm^2^, p = 0.03) and 1:5, (N = 6, mean=1.03 mm^2^, p = 0.04). D. Quantification of graft perimeter. Grafts with the 1:4 dilution (N=7, mean=7.66 mm), shared a significantly longer border with the host tissue than the 1:3 (N = 6, mean=4.36 mm, p= 0.03) and 1:5mm^2^ (N = 6, mean=3.92 mm, p = 00.02). E. Quantification of vessel density in grafts. Contralateral cortex had significantly higher vessel density than grafts with the 1:3 dilution (N=3, mean= 11.72 mm^2^, p=0.02) and 1:4 dilution (N=3, mean=8.97 mm^2^, p=0.002). Grafts with the 1:5 dilution (N=4, mean=14.18 mm^2^, p=0.13) were not statistically different than contralateral cortex.

### Donor neural precursor cells survive and differentiate into several cortical cell types

To begin assessing whether the lesion and scaffold combination supports donor cell engraftment, we transplanted dissociated mouse cortical cells harvested from embryonic day (E)12.5 telencephalon. The E12.5 telencephalon contains the cell types needed for proper development, such as neuronal, vascular, and glial precursor cells, and therefore is a practical source of cells for a proof of concept for this transplant model (Fig. 1A) (Vivian S. Chen et al., 2017). To trace donor cells, we harvested E12.5 telencephalon from *Foxg1^cre/+^;Rosa26^CAG-loxSTOPlox-eGFP/+^* mice, in which telencephalic cells are specifically labeled with GFP. Immediately after tissue dissociation, the donor cells were resuspended in the Matrigel;media solution with the addition of methylprednisolone, a corticosteroid known to decrease acute inflammation specifically in the context of brain transplantations (Liaudanskaya et al., 2019). Once the lesion was generated, the site was rinsed with cold PBS until any bleeding ceased (usually 5-20 minutes). The cell/scaffold solution was then used to fill the lesion site, and the solution gelled within 10-30 minutes. At 2 weeks post transplantation (wpt), we assessed the cell type composition of the grafts and found that GFP+ cell bodies filled the entire space in the lesion and were not observed outside of the graft site (Fig. 1B).

**Figure 1.**
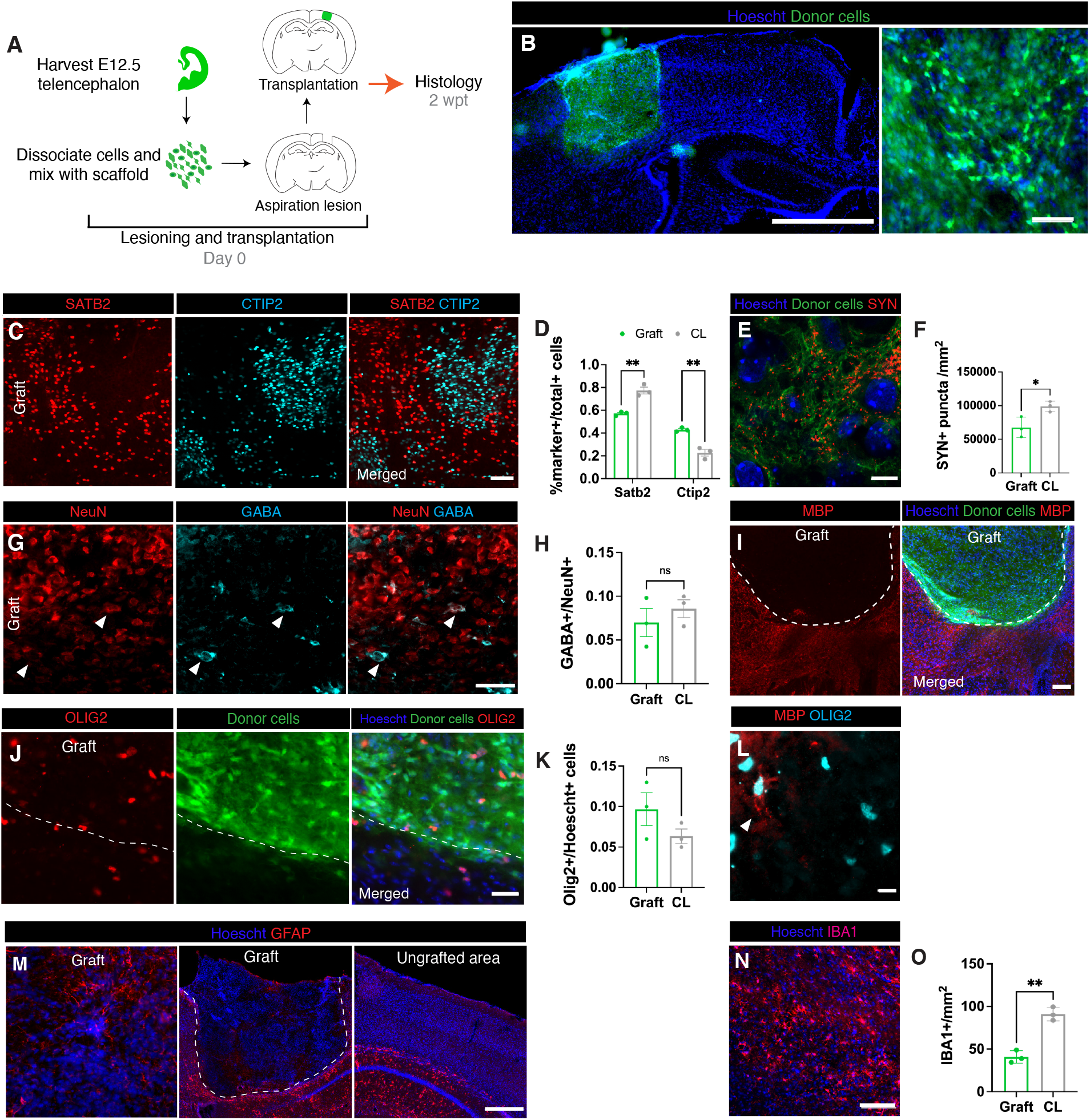
Transplanted embryonic cells in matrix differentiate at the site of aspiration lesions. A. Experimental design. B. Representative immunofluorescence image of transplant at 2 wpt at low (Left; Scale bar=1550 μM) and high (Right; Scale bar=100 μM) magnification. C. Representative immunofluorescence image of upper (SATB2) and deeper (CTIP2) cortical neurons in a graft (Scale bar=100 μM). D. Proportion of cells positive for cortical layer markers (SATB2: graft 57% of total SATB2 and CTIP2-labeled cells, N = 3; contralateral cortex 77%, N = 3, p = 0.003; CTIP2: graft 43%, N = 3; contralateral cortex 23% p = 0.003). E. Representative immunofluorescence image of anti-SYN staining (synapses) in a graft (Scale bar=5 μM). F. Density of anti-SYN fluorescence (graft 67,610 SYN+ puncta/mm^2^, N = 3; contralateral cortex 98,886 SYN+ puncta/mm^2^, N = 3, p = 0.03). G. Representative immunofluorescence image of mature neurons and inhibitory neurons (Scale bar=50 μM). Triangles indicate NeuN+/GABA+ cells. H. Quantification of inhibitory neurons (GABA) out of total mature neurons (NeuN) (graft: 7% GABA+ out of total NeuN+ cells, N=3; contralateral cortex, 9% GABA+, N = 3, p = 0.45). I. Representative immunofluorescence image of myelin along the graft host border (dotted line) (Scale bar=200 μM). J. Representative immunofluorescence image of OLIG2+ oligodendrocyte lineage cells along the graft host border (dotted line) (Scale bar= 50 μM). K. Quantification of OLIG2+/Hoescht+ cells (graft 9.7%, N = 3, contralateral cortex 6.3% N = 3, p = 0.2). L. Representative immunofluorescence image of a rare myelin (MBP)-positive oligodendrocyte in a graft (Scale = 20 μM). Triangle indicates a myelinated OLIG2+ cell. M. Representative immunofluorescence images of astrocytes in a graft and contralateral cortex. Dotted line indicates outline of graft (Scale bar=500 μM). N. Representative immunofluorescence image of IBA1+ cells in a graft (Scale bar=100 μM). O. Density of IBA1+ cells (graft 40.8/mm2 N = 3, contralateral cortex 91.1/ mm2, p = 0.21).

The neuronal donor cells differentiated into upper layer SATB2+ and deep layer CTIP2+ neurons (Fig. 1C). The ratio of upper to deep layer neurons is crucial for function (Fang et al., 2014), and SATB2+ cells are normally more abundant (Fig. 1D). We observed that at 2 wpt an abnormally low proportion of SATB2+ cells were present in the graft compared to CTIP2+ (SATB2: graft 57% of total SATB2+CTIP2+ cells, N = 3; contralateral cortex 77%, N = 3, p = 0.003; CTIP2: graft 43%, N = 3; contralateral cortex 23% p = 0.003) (Fig. 1D). The neurons in the graft also expressed synaptophysin (Fig. 1E-F), a transmembrane protein that is part of synaptic vesicles often used as a marker for presynaptic terminals (Wiedenmann & Franke, 1985). The density of synaptophysin puncta in the graft was lower than the contralateral density (graft 67,610 SYN+ puncta/mm^2^, N = 3; contralateral cortex 98,866 SYN+ puncta/mm^2^, N = 3, p = 0.03).

The cortex, among other brain regions, requires a balance of excitatory and inhibitory neurons in order to maintain and modulate information flow (Williams & Riedemann, 2021). Interneurons (GABA+) and excitatory neurons (GABA-/NeuN+) were both present in the graft at levels comparable to the contralateral cortex (graft: 7% GABA+ out of total NeuN+ cells, N=3; contralateral cortex, 9% GABA+, N = 3, p = 0.45) (Fig. 1G-H).

Glial cells including oligodendrocytes, astrocytes, and microglia make up a large part of the brain and are required for brain development, maintenance, and function (Jäkel & Dimou, 2017). Mature oligodendrocytes produce myelin that insulates axons to facilitate rapid signal transmission. Cells in the oligodendrocyte lineage (OLIG2+) were present in the grafts at a similar density to the contralateral cortex (graft 9.7%, N = 3, contralateral cortex 6.3% N = 3, p = 0.2; Fig 1J-K). However, staining for myelin basic protein (MBP) revealed that grafts at 2 wpt were largely unmyelinated (Fig. 1I). Some of the OLIG2+ cells had thin MBP+ processes extending from the cell body (Fig. 1L), suggesting that the oligodendrocytes in the grafts were immature and might produce myelin at later timepoints.

Astrocytes are crucial for neuronal maturation and function and are an essential part of the neurovascular unit which forms the blood brain barrier (BBB) (Clarke & Barres, 2013; Sweeney et al., 2019). GFAP is a marker for quiescent astrocytes, but it is not normally detected quiescent astrocytes in the neocortical parenchyma above the corpus callosum (Hinkle et al., 1997). Nevertheless, GFAP+ astrocytes were detectable in a few peripheral areas of the graft (Fig. 1M), suggesting lingering inflammation. However, given the unevenness and sparsity of these areas, GFAP+ cells were not quantified.

Microglia are essential to central nervous system development and homeostasis, playing roles in processes such as synaptic pruning, brain vascularization, and BBB maintenance (Arnold & Betsholtz, 2013; da Fonseca et al., 2014; Lenz & Nelson, 2018). Therefore, microglia are likely vital to the maturation and function of grafted cortical-like tissue. As expected, microglia (IBA1+) were present in the grafts at 2 wpt (Fig. 1N-0), albeit at a lower density than contralateral cortex (graft 40.8 cells/mm^2^ N = 3, contralateral cortex 91.1/mm^2^, p = 0.21). Microglia migrate into the mouse cortex from E9.5 until E14.5, and are therefore present in our donor cell mix (Krzyspiak et al., 2022). However, because microglia do not express *Foxg1* and are thus not labeled with GFP, microglia in the graft could be donor- or host-derived.

### Transplants become vascularized with vessels and perfused with blood

To determine whether the grafts become vascularized using the same experimental setup as in Figure 1A, we co-immunostained grafts with the vascular endothelial markers CD31 and CD105. At 2 wpt, vessels appear present throughout the grafts (Fig. 2A-B). Vascularization is a dynamic process that occurs in response to hypoxic cues in the environment, therefore we examined the graft vessels at a later time point (Andreone et al., 2015). The vessels in the graft appeared to still be changing their morphology changes until at least 4 wpt (Fig. 2C-E). From 2 to 4 wpt, despite having similar densities (2 wpt 12.2% vessels % area, N = 6; 4 wpt 8.3% vessel % area, N = 3, p = 0.1) and numbers of junctions at 2 and 4 wpt (2 wpt 79/mm^2^, N = 6; 4 wpt 30.1/mm^2^, N = 3, p = 0.08), the grafts exhibited significantly increased lacunarity at 4 wpt, a measure of irregularity or gaps within the network that does not rely on vessel density or caliber (Gould et al., 2011) (2 wpt 0.72, N = 6; 4 wpt 1.1, N = 3, p = 0.02). Consistent with ongoing maturation of graft vasculature over time, flattened Z-stacks acquired from 2-photon live imaging at 1, 2, and 4 wpt suggested that vessel length also increased between 1 and 4 wpt (Fig. 2I-J).

**Figure 2.**
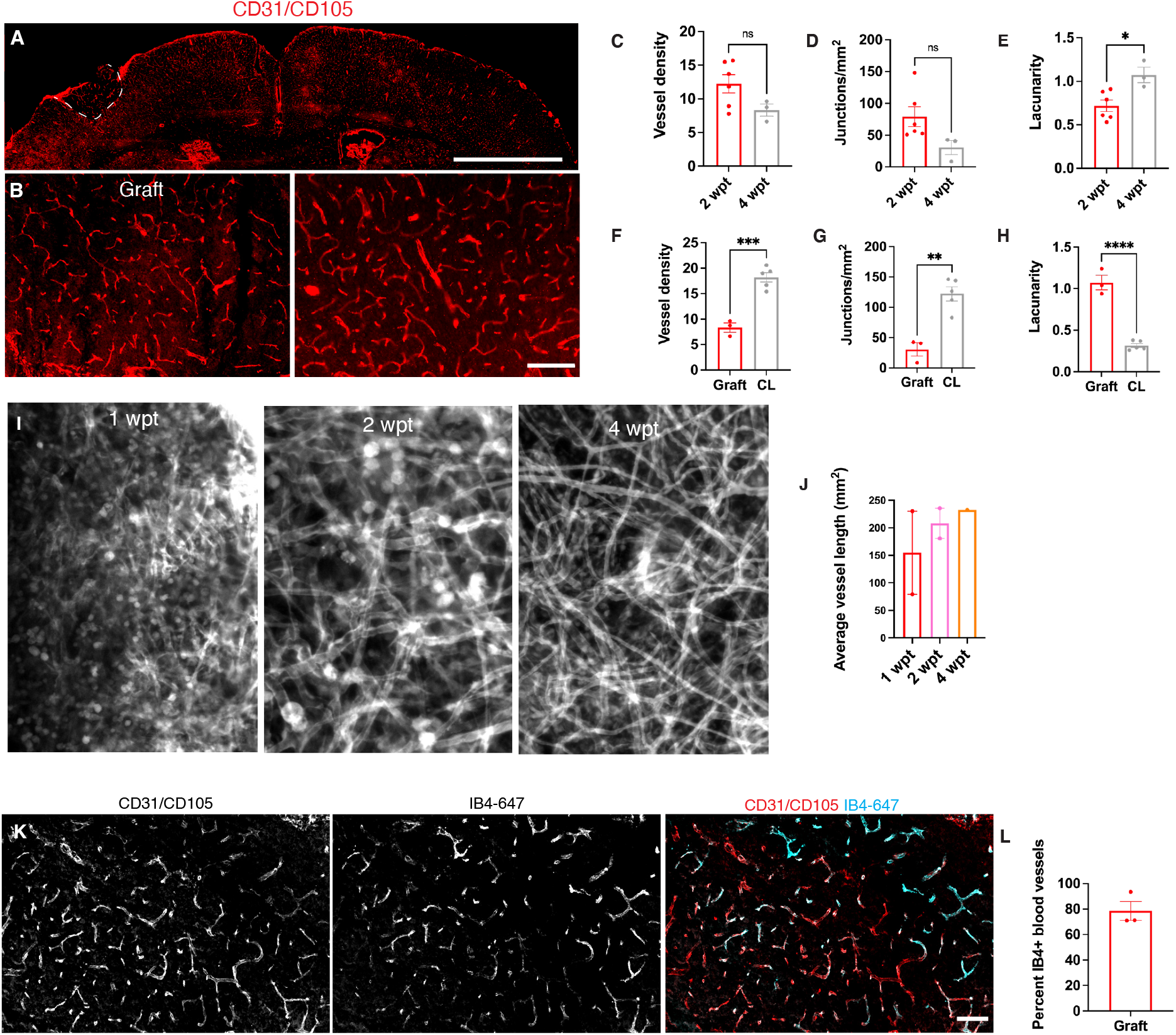
Grafts become functionally vascularized. A. Representative immunofluorescence image of vascularized cortex with graft (dotted line) at 2 wpt (Scale bar=2 mm). B. High magnification images of graft and contralateral cortex (Scale bar=200 μM). C-E. Analysis of vasculature between grafts at 2 and 4 wpt. C. Blood vessel density (2 wpt 12.2% vessel area over total area, N = 6; 4 wpt 8.3%, N = 3, p = 0.1). D. Junction and branch point density (2 wpt, 79/mm^2^, N = 6; 4 wpt, 30.1/mm^2^, N = 3, p = 0.08). E. Lacunarity (2 wpt, 0.72, N = 6; 4 wpt, 1.1, N = 3, p = 0.02). F-H. Analysis of vasculature between 4 wpt grafts and contralateral cortex. F. Blood vessel density (graft, 8.3% vessel area over total area, N = 3; contralateral cortex 18.2 %, N = 5, p = 0.0004). G. Junction and branch point density (graft 30.6/mm^2^, N = 3; contralateral cortex 122.4 N = 5, p = 0.02). H. Lacunarity (graft lacunarity = 1.1, N =3; contralateral cortex 0.3, N = 5, P < 0.0001). I. Representative 2-photon images of flattened Z-stacks of grafts at 1, 2, and 4 wpt. J. Average vessel length (1 wpt 154.2 mm^2^, N = 2; 2 wpt 208.2 mm^2^, N=2; 4 wpt 232.4 mm^2^, N = 1) K. Representative immunofluorescence image after systemic perfusion with IB4 in a graft co-stained with total vessels (Scale bar=200 μM). L. Percentage of IB4+ vessels in grafts normalized to total vessels (78.5 % IB4+ vessels, N = 3).

Since the graft vasculature appeared to still be changing within 4 wpt, we compared the graft vasculature at 4 wpt to host vasculature in the contralateral hemisphere (Fig. 2F-H). The contralateral vessels were denser than those within the graft (graft 8.3 vessel % area, N = 3; contralateral cortex 18.2 vessels % area, N = 5, p = 0.0004) and had a greater density of junctions (graft 30.6/mm^2^, N = 3; contralateral cortex 122.4%, N = 5, p = 0.02). However, the vasculature of the graft was significantly more irregular than the contralateral vasculature (graft lacunarity = 1.1, N =3; contralateral cortex 0.3, N = 5, P < 0.0001).

Vascular endothelial cells are present in mouse telencephalon at E12.5 and therefore present in the donor cell population (Krzyspiak et al., 2022) To determine the source of donor versus host vessels, we transplanted donor cells from Mesp1^cre/+^;Rosa26^loxSTOPlox-tdTomato/+^ embryos to visualize the donor vessel contribution. At 4 wpt, there were very few donor vessels found in the graft.

To test whether the graft vessels perfused blood, we intravenously injected red blood cells (RBCs) labeled with DiO and DiI and imaged blood vessel perfusion in real time with 2-photon microscopy. Blood vessel perfusion with labeled RBCs was observed as early as 7 days post-transplant (Supplementary Video 1). All of the grafts (5/5) that were imaged had vessels and RBCs circulating through them. To quantify how many of the graft vessels were perfused with blood, the mice were given intravenous injections of isolectin-B4 conjugated to Alexa Fluor 647 to label vessels through circulating blood (Fig. 2J-L). Most of the vessels in the graft were labeled with IB4-647 at 30 dpt (78.5 % IB4+ vessels, N = 3).

### Donor cells can be layered in lesion sites

Lamination is a canonical feature of the neocortex throughout development and in its mature state. For example, during mid corticogenesis, the cortex is layered into a ventricular zone, subventricular zone, subplate, cortical plate, and marginal zone. Thus, to better recapitulate the developing cortex, a transplant model should allow for the layering of cells. To test whether donor cells can be layered in a lesion by sequential gelling of scaffold-cell mixes, donor cells labeled with GFP were manually deposited in the lower half of the lesion, allowed to gel, and overlaid with donor cells labeled with tdTomato to fill the remaining space in the lesion (Fig. 3A). At 2 wpt, the donor cells were still organized in 2 layers, as originally grafted (Fig. 3B). The donor cells were not seen outside of their respective layer. Despite the distinct border between layers, cellular structures such as neural processes and blood vessels were still able to cross between the layers (Fig. 3C-D).

**Figure 3.**
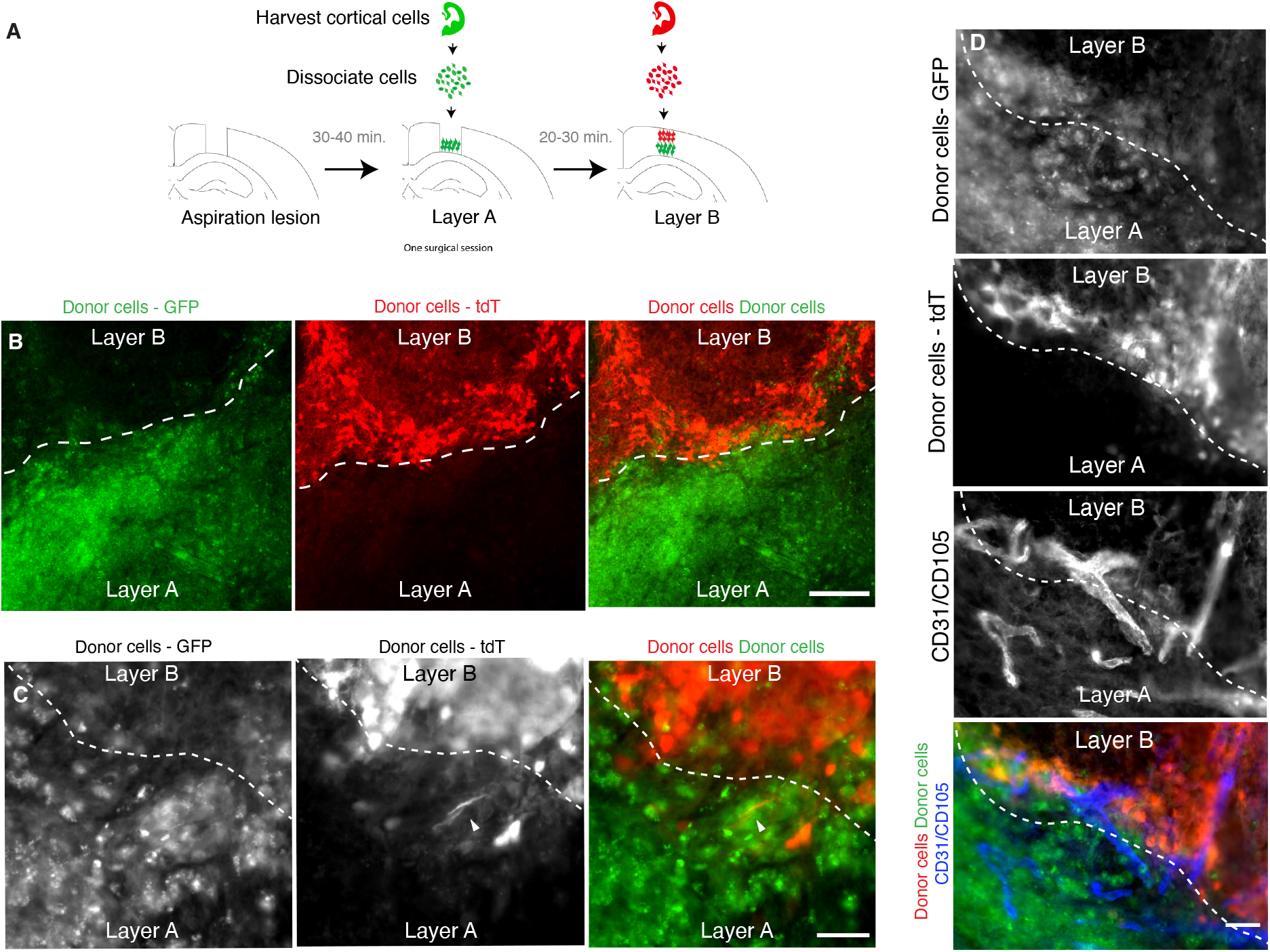
Grafts can be constructed in layers. A. Experimental design. B. Representative immunofluorescence image of a layered transplant at 2 wpt (Scale bar=200 μM). Dotted line indicates border between the layers. C. Example of a neuronal projection (arrowhead) crossing the border between layers (Scale bar=50 μM) already at 2 wpt. D. Example of blood vesselslthat cross the border between layers (Scale bar=50μM).

### Donor neurons project to appropriate targets in host brains

To be useful, a transplant model must also allow axons of donor neurons to project outside of the graft to appropriate brain targets in the host. To facilitate visualization of the donor cell processes, we transplanted cortical fetal cells that were expressing either channelrhodopsin-YFP or tdTomato into a lesion in the somatosensory cortex. At 2 wpt, the donor neurons were observed to project to several structures in the host brain (N = 7/9 mice) (Fig. 4A). Large numbers of fluorescent processes were seen projecting from the graft through the corpus callosum to the contralateral cortex (N = 6/9 mice) (Fig. 4-1,4-3,4-4) (Fenlon et al., 2017). Many fibers were present in the adjacent cortex (N=9/9) (Fig.4-2). The grafted cells projected to the striatum (N = 3/9 mice) and the hippocampus (N = 2/9 mice), known targets of the somatosensory cortex (Fig. 4-5, 4-8) (Rajasethupathy et al., 2015; Zakiewicz et al., 2014). The most ventral donor axons observed were at the lateral amygdala (N = 1/9 mice), another known target of the somatosensory cortex (Fig.4-7) (Mcdonald, 1998). The donor axons projected at least as far as 1 mm (the furthest distance examined) in both anterior (N = 2/9 mice) and posterior directions (N = 2/9 mice) at this 2 wpt timepoint (Fig4-5, 4-6). We did not observe projections in areas the axons were not expected, such as the thalamus (Fig. 4-9). Overall, this demonstrates that donor neurons reached appropriate targets already within 2 wpt in the mature host cortex (Fig. 1).

**Figure 4.**
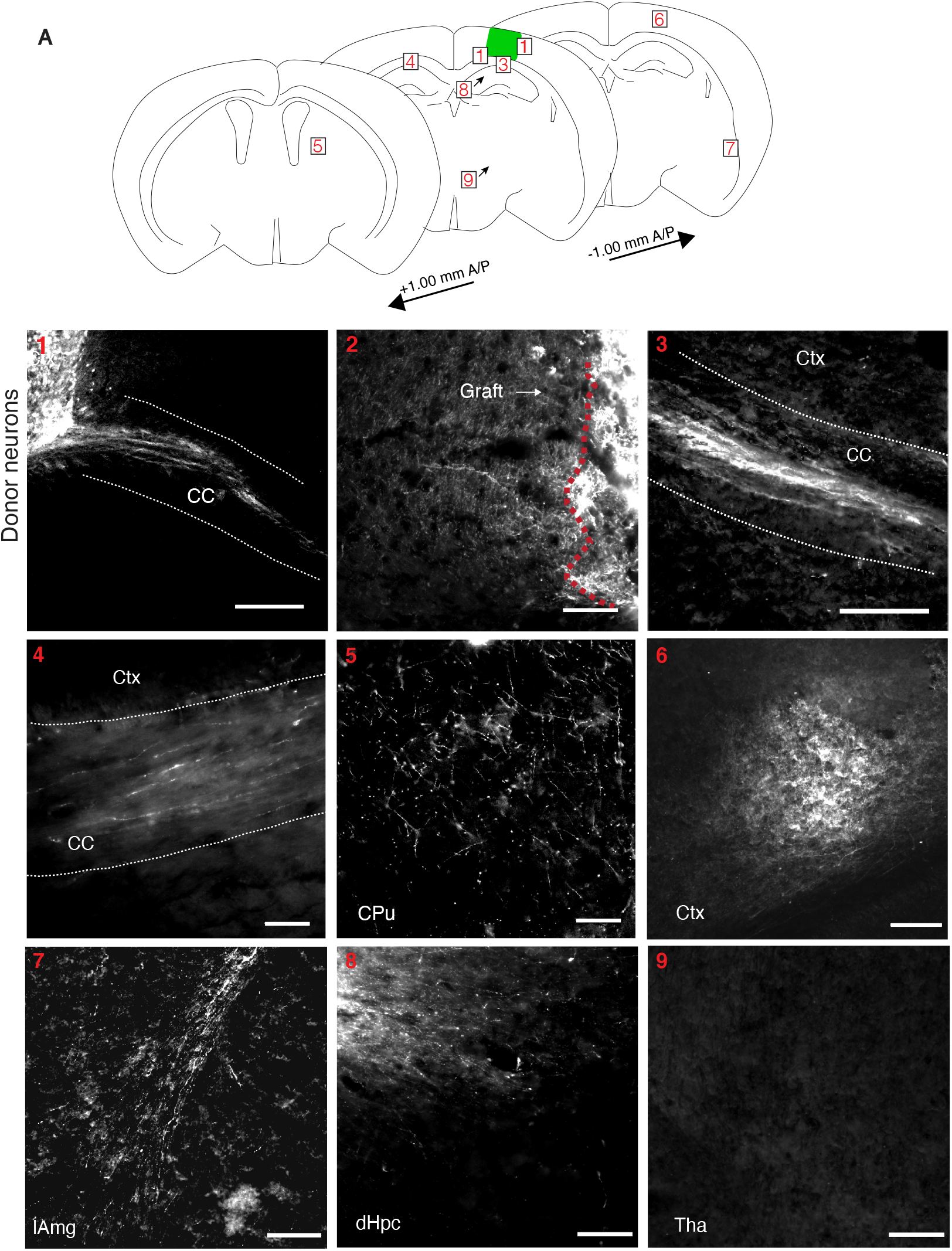
Mouse donor cells project to appropriate brain regions. A. Diagram of areas in host brain that donor cells project to at 2 wpt. Projections are seen in 1. the corpus callosum (CC) exiting the graft (N=6/9) (Scale bar=300 μM). 2. the cortex directly adjacent to the graft (N=9/9) (Scale bar=80 μM). 3. the CC away from the graft towards the contralateral hemisphere (N=6/9) (Scale bar=80 μM). 4. the CC in contralateral cortex (N=6/9) (Scale bar=40 μM). 5. the caudate putamen (CPu) (N=3/9) (Scale bar=80 μM). 6. the posterior cortex (N=2/9) (Scale bar=100 μM). 7. the lateral amygdala (lAmg) (N=1/9) (Scale bar=200 μM). 8. the dorsal hippocampus (dHPC) (N=2/9) (Scale bar=100 μM). 9. and the thalamus (Thal) (N=9/9) (Scale bar=100 μM).

### Transplanted neurons become electrophysiologically active and respond to visual stimuli

To test whether the donor neurons functionally integrated in the cortical network and were able to fire action potentials, integrate with the host, and respond to external inputs such as sensory stimuli, we transplanted cells into the visual cortex (V1). An ultra-flexible neural probe designed for long-term recordings was inserted in the scaffold-cell mix at the time of transplantation and into the contralateral V1 that was intact and served as a control (Fig. 5A) (Yin et al., 2022). First, to observe the maturation of the donor neurons, we recorded non-evoked activity from freely moving mice starting 2-3 days after transplantation for 30 minute sessions, 3 times a week for 12 weeks. At 2 wpt there was very little activity. By 3 wpt, there was an increase of spikes and local field potential (LFP) consistent with the emergence and increase of synaptogenesis (Fig. 5B). One month after transplantation the “noise” subsided and the magnitude of the action potentials increased, possibly indicative of pruned synapses and maturing cortical tissue. Finally, at 5 wpt, higher amplitude spikes at a more regular frequency were observed, consistent with stable network activity. At 9 and 13 wpt, graft local field potential (LFP) power spectra are similar to the control. The corresponding LFP traces confirm the power spectra. Thus, the grafted neurons in this transplant model were maturing along a normal schedule and fired spontaneous action potentials (Luhmann et al., 2016).

**Figure 5.**
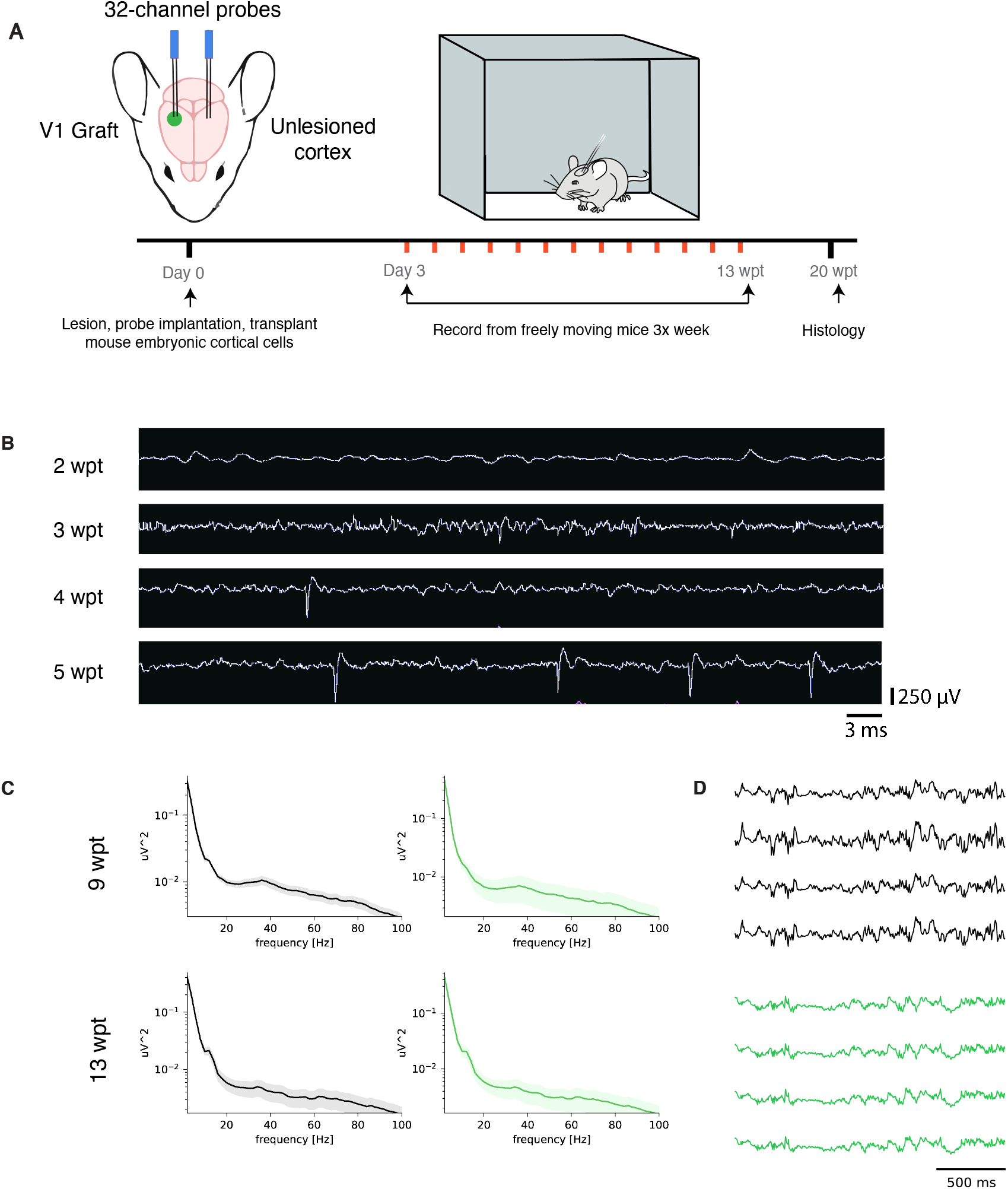
Mouse donor neurons become physiologically active. A. Experimental design. B. Neural traces of a single representative channel at 2, 3, 4, and 5 wpt with the same time and voltage scales. C. LFP power spectrums of control (left panels) and graft (right panels) at 9 and 13 wpt. D. LFP traces of control (gray) and graft (green) at 9 and 13 wpt.

To determine whether the donor neurons integrated into the visual cortical network and responded to external visual stimuli, we recorded from mice that were head-fixed and presented visual stimuli once a week from 1-13 wpt (Fig. 6A). The visual stimulus program displayed bars in 6 orientations and moving in 2 directions (right to left and left to right). Recorded units were classified as regular spiking cells (RS), which are putative excitatory neurons, and fast spiking cells (FS), which are putative inhibitory interneurons. Examples of spike waveforms of FS corresponding to putative excitatory neurons in the control (Spike width = 1.05 ms, FR = 3.2 HZ) and graft (Spike width = 1.10 ms, FR = 2.3 HZ) are shown in Figure 6B. Neurons in the graft became visually responsive (N = 2/3) as early at 2 wpt (SNR = 0.175, p = 0.002), however at this early transplant ages the responses of donor neurons were predominantly present at stimuls onset(Fig 6C). At 4 wpt, there was evidence of neuronal maturation. For example, a putative donor neuron displayed increased signal to noise ratio (SNR = 0.303, p < 0.001) and fired at stimulus onset, consistent with maturation. However, not all neurons appeared to be maturing at the same rate since a different at 4 wpt continued to display a stimulus offset response (1 second delay) with a lower snr (SNR = 0.234, p = 0.001) (right panel). At 8 wpt, the firing rate was topically increased at stimulus onset and sustained for 1 sec, followed by 1.5 seconds of low firing rate (SNR = 0.919, p < 0.001). Finally, at 13 wpt we consistently observed SNR comparable to those of units in the control side (SNR = 0.0845, p < 0.001) which included donor neurons firing sparsely and being highly driven by stimulus onset, consistent with donor cells maturing into adult functional pyramidal neuron. The orientation selectivity index (OSI) of the donor neurons increased between 2 (OSI = 0.23) and 4 wpt (OSI = 0.25 (Fig. 6D), indicating the graft was becoming more tuned to specific orientations. These data provide evidence that the donor neurons in primary visual cortex are synaptically integrating with the host and processing external visual stimuli, potentially via thalamic input, as would normal developing V1 cortex (Shen & Colonnese, 2016).

**Figure 6.**
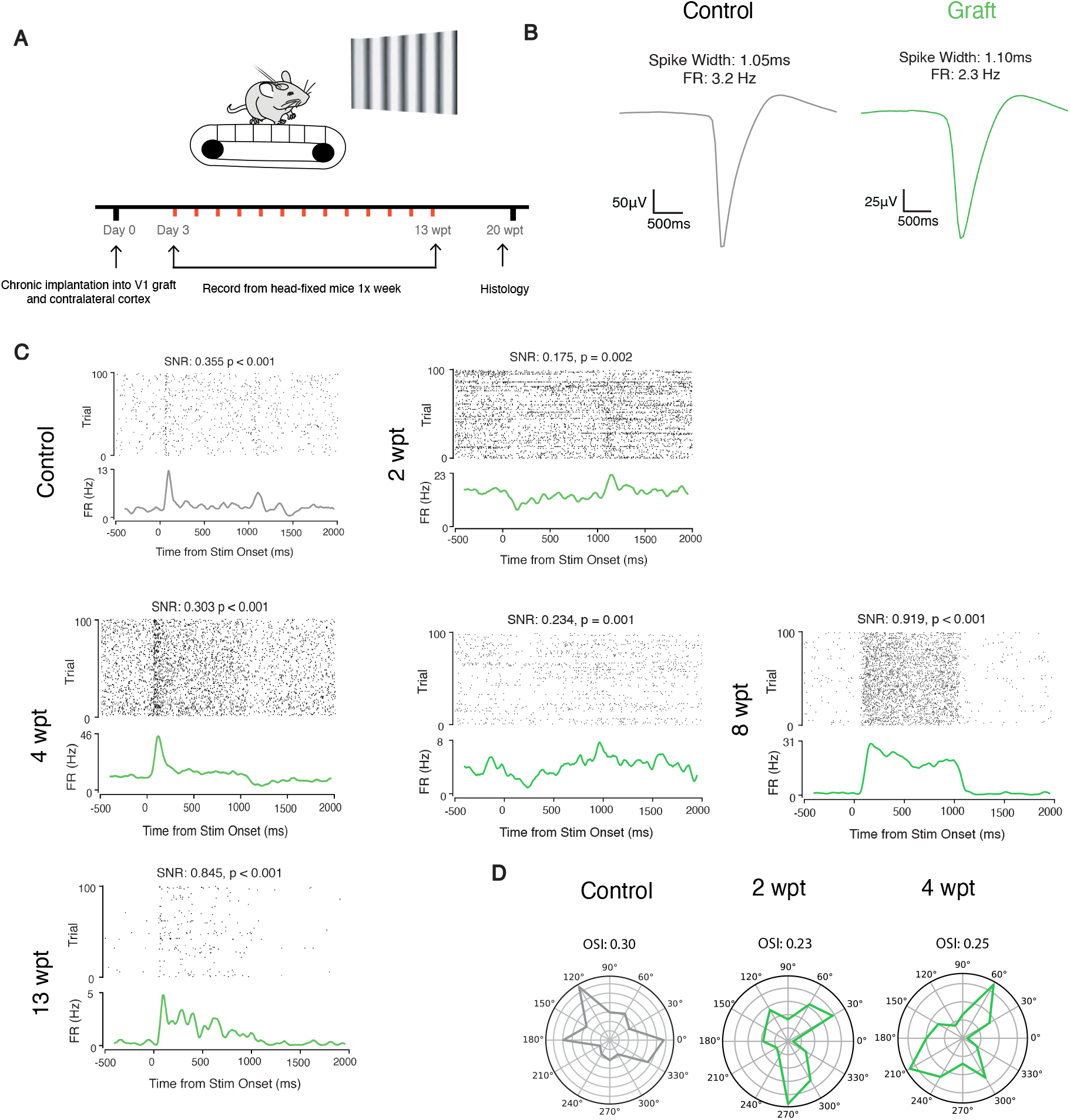
Mouse donor neurons respond to visual stimuli and are tuned to specific orientations. A. Experimental design. B. Representative waveforms of putative neurons from control (left) and graft (right). C. Raster plots of firing rate and SNR for control and graft at 2, 4, 8, and 13 wpt (from left to right, top to bottom) D. OSI for control and graft at 2 and 4 wpt.

## Discussion

As a platform to rebuild neocortical tissue *in vivo*, we created a cortical transplant model that can support multiple cortical cell types, is vascularized, and is amenable to layering. The majority of cortical transplantation studies rely on the injection of cells directly into the parenchyma, such as through a Hamilton syringe or pulled glass pipette (Ballout et al., 2016; Espuny-Camacho et al., 2018; Real et al., 2018). Such methods provide the experimenter with little control over the exact shape of the graft or arrangement of the donor cells within it, resulting in disorganized and unpredictable tissue structures.

To generate consistently sized lesions and increase transplant precision, we used an aspiration lesion. In this study, we removed healthy host cortical tissue, but in the future a similar approach may be used to remove degenerated or scarred tissue. The space created by the aspiration lesion then allows for the grafting of donor cells with a scaffold to provide more control over the final overall shape and size of the resulting tissue. Scaffolds and signaling factors within them can also increase donor cell survival and differentiation (Carlson et al., 2016; Wang et al., 2020).

In addition, our transplant model provides a platform for organizing cells into layers. The mature neocortex has a distinct 6-layered structure, with each layer containing specific combinations of neuronal subtypes. While cortical transplantation studies show that subtypes of neurons differentiate in grafts, they are not laminated as in the mature cortex (Agirman et al., 2017; Palma-Tortosa et al., 2020). Lack of normal lamination could prevent proper wiring of the donor neurons both with each other and with the host cortex. The method described here allows for the layering of precursor cells, which should in the future provide better control in achieving mature tissue that more closely recapitulates normal neocortex. For example, the layering of the different cell types that make up the developing ventricular zone, subventricular zone, subplate, and marginal zone (Agirman et al., 2017), could increase the likelihood of normal development of the transplant and better integration into the host cortex, potentially allowing for true cortical repair.

Another benefit of the grafting platform described here is that new tissue is assembled *in vivo*, which facilitates its rapid vascularization. Nutrients and oxygen can only diffuse ~400 um, necessitating any aerobic tissue larger than 400 um to establish and maintain a vascular system (Auger et al., 2013). Vascularization is often a major hurdle in the expansion and long-term survival of 3D cultures and bioengineered tissues (Auger et al., 2013). Proper vascularization of brain tissue is more of a challenge because it is highly metabolic, requiring 20% of the total oxygen consumed by a human (Hawkins & Davis, 2005). Vessels in the graft are observed as early as 2 dpt, with significant vascularization occurring by 7 dpt. Neurogenic niches in mammals have lower levels of oxygen compared to more mature tissue, so the short delay in vascularization immediately post-transplant may not be an issue (De Filippis & Delia, 2011).

Despite the presence of vascular endothelial cells in our donor cell population and their importance in contributing new functional blood vessels after transplantation at the site of ischemic strokes (Krzyspiak et al., 2022), the vessels observed here after transplantation to the sites of aspiration lesions are primarily host-derived. Nevertheless, we show that the vessels in these grafts have circulating RBCs and are therefore likely transporting oxygen and nutrients to the new tissue to promote survival and integration.

Importantly, our methods produces neurons that integrate into functional networks and are spontaneously active driven by sensory inputs, This demonstrates that our transplant model is suitable for functional neurons.

Current human disease modeling mostly relies on *in vitro* models derived from human pluripotent stem cells, which are often simple in their cell type composition and variable in their structure, in addition to being constrained by short survival due to the absence of vascularization and blood circulation (Giandomenico & Lancaster, 2017). Therefore, *in vivo* models to study human tissues would be advantageous. Interspecies chimeric models are becoming increasingly accepted as a method to investigate human diseases (Espuny-Camacho et al., 2013; Real et al., 2018). Our study suggests that the adult neocortex of immune compromised mice could be used as a platform for studying the cellular and molecular interactions between human neurons, vascular cells, and glia, and serve as a model for studying human neocortical disease, drug testing, and therapeutic tissue replacement.

## Methods

### Animals

Animal experiments were approved by the Albert Einstein College of Medicine Institutional Animal Care and Use Committee in accordance with National Institutes of Health guidelines. Swiss Webster mice (Charles River), male and female, ages 2-6 months were used as hosts for this study. Donor embryos were harvested from transgenic mice appropriate for the experiment. For visualizing neural components, donors were from *Foxg1^cre/+^* (Jax # 06084) mice crossed with homozygous *Rosa26^CAG-loxSTOPlox-eGFP^* (Jax #010701) mice. For visualizing graft-derived vascularization, donors were from Mesp1^cre/+^ mice (Saga et al., 1999) (donated from Paul Frenette’s lab, Albert Einstein College of Medicine) crossed with homozygous Rosa26^loxSTOPlox-tdTomato^ mice (Jax # 007909). For visualizing graft layers, donors were from *Foxg1^cre/+^* mice crossed with homozygous *Rosa26^CAG-loxSTOPlox-eGFP^* mice, and donors from *Foxg1^cre/+^* mice crossed with homozygous Rosa26^loxSTOPlox-tdTomato^ mice. For graft projections, donors were from *Foxg1^cre/+^* mice crossed with homozygous *Rosa26^CAG-loxSTOPlox-Chr2-EYFP^* (Jax # 012569) mice, or *Foxg1^cre/+^* mice crossed with homozygous Rosa26^loxSTOPlox-tdTomato^. Mice were housed on a 12 h light-dark cycle and provided with water and chow ad libitum.

### Fetal telencephalon dissociation

Donor cells were harvested and dissociated from E12.5 telencephalons with genotypes suitable for each experiment (see “Animals” above). Dissociation was performed with Accutase (Innovative cell technologies, California, United States, cat. AT104) followed by trituration of the cell suspension. Cells were washed, counted, and then resuspended at ~500k cells per ul in mouse embryonic cell media (Neurobasal, B27, N2, pen/strep) diluted with Matrigel (Corning, New York, United States, cat. 356234), with the addition of Methylprednisolone (8.7mM; Sigma, Missouri, United States, cat. M3781) and VEGF (20ng/ml; Sigma cat. GF445).

### Transplantation procedures

Mice were placed under 5% isoflurane infused oxygen anesthesia and once anesthetized maintained at 2% isoflurane/oxygen. Mice were placed on a stereotaxic platform and head fixed with ear bars. The scalp was wiped with betadine prior to making a 6 mm incision over the midline. A 2mm diameter craniotomy was made over the somatosensory cortex (centered at Bregma: A/P ~1.0, M/L ~2.0) followed by an incision in the dura with a scalpel. The dura was carefully peeled back and a biopsy punch was inserted 1 mm deep into the cortex. Any tissue within the cut area that was not extracted with the biopsy punch was removed with a blunt tip needle attached to a vacuum to create a uniform cylindrical lesion. The blood was flushed with saline until the bleeding stopped. The cell/Matrigel solution was then added to fill the lesion. Once the cell solution had gelled, a 3mm cover glass window was superglued to seal the craniotomy. The scalp was sutured shut and the mouse was given subcutaneous injections of Flunixin (FlunixiJect, Henry Schein, New York, United States) and Ceftriaxone (Pfizer, Massachusetts, United States) and monitored closely for 24 h followed by daily monitoring for signs of infection or morbidity. In cases where layering of cells was performed, the procedure was similar except GFP+ cells were added to fill half of the lesion (about 0.75 ul) to generate layer A. Once layer A had gelled (approximately 20-30 min.) tdT+ cells were added to the lesion on top of the GFP+ cells to form layer B.

### Tissue processing and Immunohistochemistry

Mice were anesthetized and then perfused with saline followed by 4% paraformaldehyde in PBS (PFA/PBS), via IV cardiac puncture. Brains were removed, fixed overnight in PFA/PBS, placed in 30% sucrose for 1-2 days, frozen in OCT, and cryosectioned at 30 μm. Sections were incubated in blocking buffer (10% normal goat serum in 0.3% triton in PBS) for 1 h, incubated with primary antibodies (see antibody list) overnight at 4°C, washed with PBS, incubated with secondary antibody (1:600) and Hoescht (1:10,000) for 1-2 h at RT, washed, mounted with Flouromount G (ThermoFisher, New Jersey, United States, cat. 00-4958-02), and topped with a glass cover. Stained sections were imaged with an epifluorescent microscope.

### Image analysis

To quantify cortical markers, interneurons, oligodendrocyte lineage-cells, and microglia in the graft, cells from 3 sections per mouse were counted by hand in Photoshop. For the contralateral cortex, the entire dorsal/ventral length from the pia to the corpus callosum was selected for quantification to account for variation in density between layers. To quantify synapses, SYN+ puncta was counted by hand. The software Angiotool was used to analyze the blood vessels (Zudaire et al., 2011). The flattened Z-stack of graft vasculature was generated with Image J(Schneider et al., 2012). To quantify the amount of vessels that are being perfused with blood, IB4+ vessels were divided by the total amount of vessels.

### *In vivo* live imaging

Mice were imaged with a two-photon laser scanning microscope (Thorlabs Bergamo) using a femtosecond-pulsed laser (Chameleon Ultra II, Coherent) tuned to 910-nm or a 1,055-nm femtosecond-pulsed laser (Fidelity 2, Coherent) and a 16X water immersion objective (0.8NA, Nikon, New York, United States). ThorImage software was used to acquire the images. The mice were anesthetized with 1-2% isoflurane and head-fixed. To visualize blood vessel structure, Z-stacks starting from the dorsal side of the graft were taken.

### Perfusion of blood vessels

To visualize perfusion, mice were injected intravenously with labeled RBCs. Briefly, after harvesting blood from mice, RBCs were segregated from plasma and serum using centrifugation (500g, 5 min), washed with saline, incubated for 1 hr at 37°C with DiI (1:50) and DiO (1:50) (ThermoFisher cat. V2285; V2286), washed, and resuspended at 50% hematocrit. 100 ul of DiO/DiI-labeled RBCs were injected through the retro-orbital sinus while mice were under anesthesia. Labeled RBCs could be seen in the vessels up to 3 weeks post injection. To label blood vessels post-hoc, 100 μL of IB4-647 was injected retro-orbitally 30 minutes before euthanasia.

### Headpost and chronic electrode implantation surgery

On the day of the surgery, the mouse was anesthetized with isoflurane and the scalp was shaved and cleaned three times with Betadine solution. An incision was made at the midline and the scalp resected to each side to leave an open area of skull. A head-post was glued with dental cement to the skull. Two 3 mm craniotomies were made on both hemispheres centered at Bregma: A/P −2.7, M/L 2.5. On one hemisphere, a lesion was made in V1 followed by the transplantation of donor cells from Foxg1^cre/+^;Rosa26^CAG-loxSTOPlox-eGFP/+^ mice. Mice were implanted with 2 probes, one in the graft and one in contralateral intact V1. Probes had 32 channels either in a linear or tetrode arrangement. Once probes were inserted, a 3 mm glass window was applied to each craniotomy then sealed with Kwik-Sil followed by super glue. One skull screw (McMaster-Carr) was placed at posterior pole. The skin was then glued to the edge of the Metabond with cyanoacrylate. Metabond was applied to secure the probes on the head. Analgesics were given immediately after the surgery and on the two following days to aid recovery. Mice were given a course of antibiotics (Sulfatrim, Butler Schein, Florida, United States) to prevent infection and were allowed to recover for 3-5 days following implant surgery before beginning spontaneous freely moving recordings. Mice were handled for 10 min/day for 5 days prior to the headpost surgery. After recovering from implant surgery, mice were habituated to treadmill fixation for 30-60 minutes per day for a total of 5 days.

### In vivo electrophysiology

All extracellular single-unit and LFP recordings were made with chronic ultra-flexible neural probes designed for eliciting minimal inflammation and for long-term recordings (Luan et al., 2017; Yin et al., 2022). Signals were digitized and recorded by the RHD recording system (Intan, California, United States). Data were sampled at 30kHz for freely moving sessions and 20kHz for head-fixed visual stimulus sessions. Recordings were performed during the light portion of the light/dark cycle.

### Visual stimulation

Visual stimuli were presented on an LCD monitor at a spatial resolution of 1080 x1080 pixels/cm, with a real-time frame rate of 60Hz, and a mean luminance of 50 cd/m^2^ positioned 15cm from the eye. The LCD monitors used for visual stimulation were positioned on both sides of the animal, perpendicular to the surface of each eye. Animals underwent two recording sessions per day, with monocular stimulation alternating between sessions. Both control and graft sides were recorded during the sessions regardless of the eye that was stimulated. Each session contained a 20 minute block of gray screen followed by one-hundred percent contrast drifting gratings in 12 orientations with a spatial frequency of 0.2 cycles/cm and temporal frequency of 2 cycles/second. Each stimuli was presented for 1 second followed by a 1.5 second interstimulus interval with 0.5 seconds jitter.

### In vivo electrophysiology analysis

Spikes were clustered semi-automatically using the following procedure. The Matlab package Kilosort2 was used to identify clusters through template matching (Pachitariu et al., 2016). Fast and accurate spike sorting of high-channel count probes was performed with Kilosort. Well-isolated units were the curated using the Python library graphical user interface phy 2.0 (Rossant & Harris, 2013). Hardware-accelerated interactive data visualization for neuroscience in Python. Clusters contained no more than 10% auto correlogram contamination with a refractory period of 2ms. Unit data were analyzed for firing rates and visual responses using custom-written Python code.

### Single unit activity analysis in head-fixed recordings

Spike waveforms were extracted and averaged from the high-pass filtered signal around the time of peak unit amplitude detected by Kilosort2. Firing rate was computed by dividing the total number of spikes a cell fired in a specified period by the total duration of that period. Quantification of rate modulation to visual stimulus was computed as the firing rate in the 400 ms period after stimulus onset, and the firing rate in the 500 ms before stimulus onset. Signal to noise ratio (SNR) was computed as the absolute value of the stimulus-evoked firing rate minus baseline firing rate divided by the sum of the baseline firing rate and the stimulus-evoked firing rate: SNR = abs((FR_evoked - FR_baseline)/(FR_evoked + FR_baseline)). Orientation selectivity index was calculated as one minus the circular variance as described by Batista-Brito et al. (2017). Here, values larger than 0 show that the cell fired more than chance for that orientation while an OSI=1 indicates that the cell only fires for a single stimulus orientation.

### Local field potential computation in freely-moving recordings

Raw electrophysiology data were processed using SciPy signal processing toolbox. A low-pass Butterworth filter was applied at 200Hz and subsequently downsampled to 1250Hz. LFP power spectra were computed using Welch’s method, with a 500ms Hanning taper and normalized over the total estimated power of the signal. Spectra from control and graft sides were averaged across 3-4 channels in a single animal for each recording time point.

### Statistical analyses

Statistical analyses were performed using GraphPad Prism. Data were analyzed for statistical significance using unpaired t test for the quantification of interneurons, oligodendrocytes, microglia, synapses, and blood vessels in the graft compared with the contralateral control. Multiple unpaired t test was used for quantitation of cortical layer subtypes. One-way ANOVA was used for quantitation of experiments examining Matrigel dilution. The p values and sample sizes for each experiment are listed in the results section and figure legends.

**Table 1.** Antibody list. Asterisks indicate that antigen retrieval is recommended.

| Antibody | Company | Catalog no. | Species | Dilution |
| --- | --- | --- | --- | --- |
| Anti-CD105 | Biolegend | 120402 | Rat | 1:50* |
| Anti-CD31 | BD Pharmingen | 553370 | Rat | 1:50* |
| Anti-CTIP2 | abcam | ab18465 | Rat | 1:500* |
| Anti-GABA | Millipore-Sigma | ab2052 | Rabbit | 1:1000 |
| Anti-GFAP | Invitrogen | 13-0300 | Rat | 1:500 |
| Anti-GFP | ThermoFisher | A11122 | Rabbit | 1:250 |
| Hoechst 33342 | Invitrogen | H3570 | N/A | 1:10,000 |
| IB4-647 | ThermoFisher | I32450 | N/A | 1µg/µl |
| Anti-Iba1 | Wako | 019-19741 | Rabbit | 1:100 |
| Anti-MBP | abcam | ab7349 | Rat | 1:50 |
| Anti-NeuN | Synaptic Systems | 266004 | Guinea Pig | 1:500 |
| Anti-OLIG2 | Millipore-Sigma | ab9610 | Rabbit | 1:50 |
| Anti-SATB2 | abcam | ab92446 | Rabbit | 1:500* |
| Anti-Synaptophysin | abcam | ab32127 | Rabbit | 1:500* |

## Acknowledgements

We are grateful for grant support from the SENS Research Foundation (JMH), the Methuselah Foundation (JMH), the New York State Department of Health NYSTEM Program for shared facility grant support C029154 (JMH), NYSTEM C34875GG (JMH), NYSTEM Einstein Training Program in Stem Cell Research C30292GG (AQ), Brain Research Foundation (JMH), NIH NS088943 (JMH), NARSAD Young Investigator Award (RBB), Whitehall Foundation Research Grant (RBB), SFARI Bridge to Independence Award 381222 (RBB), National Institute of Child Health and Human Development F31 Predoctoral Fellowship (CW), and seed funding from the Albert Einstein College of Medicine. We are also grateful to lab members for feedback on the manuscript.

## References

Agirman, G., Broix, L., & Nguyen, L. (2017). Cerebral cortex development: an outside-in perspective. FEBS Letters, 591, 3978–3992. https://doi.org/10.1002/1873-3468.12924

Andreone, B. J., Lacoste, B., & Gu, C. (2015). Neuronal and Vascular Interactions. Annual Review of Neuroscience, 38, 25–46. https://doi.org/10.1146/annurev-neuro-071714-033835

Arnold, T., & Betsholtz, C. (2013). The importance of microglia in the development of the vasculature in the central nervous system. Vascular Cell, 5(4), 1–7. https://doi.org/10.1186/2045-824X-5-4

Auger, F. A., Gibot, L., & Lacroix, D. (2013). The Pivotal Role of Vascularization in Tissue Engineering. Annual Review of Biomedical Engineering, 15, 177–200. https://doi.org/10.1146/annurev-bioeng-071812-152428

Ballout, N., Frappé, I., Péron, S., Jaber, M., Zibara, K., & Gaillard, A. (2016). Development and Maturation of Embryonic Cortical Neurons Grafted into the Damaged Adult Motor Cortex. Frontiers in Neural Circuits, 10(August), 1–12. https://doi.org/10.3389/fncir.2016.00055

Batista-Brito, R., Vinck, M., Ferguson, K. A., Chang, J. T., Laubender, D., Lur, G., Mossner, J. M., Hernandez, V. G., Ramakrishnan, C., Deisseroth, K., Higley, M. J., & Cardin, J. A. (2017). Developmental Dysfunction of VIP Interneurons Impairs Cortical Circuits. Neuron, 95(4), 884–895.e9. https://doi.org/10.1016/j.neuron.2017.07.034

Carlson, A. L., Bennett, N. K., Francis, N. L., Halikere, A., Clarke, S., Moore, J. C., Hart, R. P., Paradiso, K., Wernig, M., Kohn, J., Pang, Z. P., & Moghe, P. V. (2016). Generation and transplantation of reprogrammed human neurons in the brain using 3D microtopographic scaffolds. Nature Communications, 7, 1–10. https://doi.org/10.1038/ncomms10862

Clarke, L. E., & Barres, B. A. (2013). Emerging roles of astrocytes in neural circuit development. Nature Reviews Neuroscience, 14(5), 311–321. https://doi.org/10.1038/nrn3484

da Fonseca, A. C. C., Matias, D., Garcia, C., Amaral, R., Geraldo, L. H., Freitas, C., & Lima, F. R. S. (2014). The impact of microglial activation on blood-brain barrier in brain diseases. Frontiers in Cellular Neuroscience, 8(362), 1–13. https://doi.org/10.3389/fncel.2014.00362

Daviaud, N., Friedel, R. H., & Zou, H. (2018). Vascularization and engraftment of transplanted human cerebral organoids in mouse cortex. ENeuro, 5(6), 1–18. https://doi.org/10.1523/ENEURO.0219-18.2018

De Filippis, L., & Delia, D. (2011). Hypoxia in the regulation of neural stem cells. Cellular and Molecular Life Sciences, 68, 2831–2844. https://doi.org/10.1007/s00018-011-0723-5

Espuny-Camacho, I., Michelsen, K. A., Gall, D., Linaro, D., Hasche, A., Bonnefont, J., Bali, C., Orduz, D., Bilheu, A., Herpoel, A., Lambert, N., Gaspard, N., Péron, S., Schiffmann, S. N., Giugliano, M., Gaillard, A., & Vanderhaeghen, P. (2013). Pyramidal Neurons Derived from Human Pluripotent Stem Cells Integrate Efficiently into Mouse Brain Circuits In Vivo. Neuron, 77(3), 440–456. https://doi.org/10.1016/j.neuron.2012.12.011

Espuny-Camacho, I., Michelsen, K. A., Linaro, D., Bilheu, A., Acosta-Verdugo, S., Herpoel, A., Giugliano, M., Gaillard, A., & Vanderhaeghen, P. (2018). Human Pluripotent Stem-Cell–Derived Cortical Neurons Integrate Functionally into the Lesioned Adult Murine Visual Cortex in an Area-Specific Way. Cell Reports, 23(9), 2732–2743. https://doi.org/10.1016/j.celrep.2018.04.094

Falkner, S., Grade, S., Dimou, L., Conzelmann, K. K., Bonhoeffer, T., Götz, M., & Hübener, M. (2016). Transplanted embryonic neurons integrate into adult neocortical circuits. Nature, 539(7628), 248–253. https://doi.org/10.1038/nature20113

Fang, W. Q., Chen, W. W., Jiang, L., Liu, K., Yung, W. H., Fu, A. K. Y., & Ip, N. Y. (2014). Overproduction of Upper-Layer Neurons in the Neocortex Leads to Autism-like Features in Mice. Cell Reports, 9, 1635–1643. https://doi.org/10.1016/j.celrep.2014.11.003

Fenlon, L. R., Suárez, R., & Richards, L. J. (2017). The anatomy, organisation and development of contralateral callosal projections of the mouse somatosensory cortex. Brain and Neuroscience Advances, 1, 1–9. https://doi.org/10.1177/2398212817694888

Giandomenico, S. L., & Lancaster, M. A. (2017). Probing human brain evolution and development in organoids. Current Opinion in Cell Biology, 44, 36–43. https://doi.org/10.1016/j.ceb.2017.01.001

Gould, D. J., Vadakkan, T. J., Poche, R. A., & Dickinson, M. E. (2011). Multifractal and Lacunarity Analysis of Microvascular Morphology and Remodeling. Microcirculation, 18(2), 136–151. https://doi.org/10.1111/j.1549-8719.2010.00075.x.Multifractal

Hawkins, B. T., & Davis, T. P. (2005). The blood-brain barrier in health and disease. Pharmacology Reviews, 57, 173–185. https://doi.org/10.1002/ana.23648

Henriques, D., Moreira, R., Schwamborn, J., Pereira de Almeida, L., & Mendonça, L. S. (2019). Successes and Hurdles in Stem Cells Application and Production for Brain Transplantation. Frontiers in Neuroscience, 13, 1–15. https://doi.org/10.3389/fnins.2019.01194

Hinkle, D. A., Baldwin, S. A., Scheff, S. W., & Wise, P. M. (1997). GFAP and S100β expression in the cortex and hippocampus in response to mild cortical contusion. Journal of Neurotrauma, 14(10), 729–738. https://doi.org/10.1089/neu.1997.14.729

Jäkel, S., & Dimou, L. (2017). Glial cells and their function in the adult brain: A journey through the history of their ablation. Frontiers in Cellular Neuroscience, 11, 1–17. https://doi.org/10.3389/fncel.2017.00024

Krzyspiak, J., Yan, J., Ghosh, H. S., Galinski, B., Lituma, P. J., Alvina, K., Quezada, A., Kee, S., Grońska-Pęski, M., Tai, Y. De, McDermott, K., Gonçalves, J. T., Zukin, R. S., Weiser, D. A., Castillo, P. E., Khodakhah, K., & Hébert, J. M. (2022). Donor-derived vasculature is required to support neocortical cell grafts after stroke. Stem Cell Research, 59. https://doi.org/10.1016/j.scr.2021.102642

Lee, J. P., Jeyakumar, M., Gonzalez, R., Takahashi, H., Lee, P. J., Baek, R. C., Clark, D., Rose, H., Fu, G., Clarke, J., McKercher, S., Meerloo, J., Muller, F. J., Park, K. I., Butters, T. D., Dwek, R. A., Schwartz, P., Tong, G., Wenger, D.,…Snyder, E. Y. (2007). Stem cells act through multiple mechanisms to benefit mice with neurodegenerative metabolic disease. Nature Medicine, 13(4), 439–447. https://doi.org/10.1038/nm1548

Lenz, K. M., & Nelson, L. H. (2018). Microglia and beyond: Innate immune cells as regulators of brain development and behavioral function. Frontiers in Immunology, 9(APR). https://doi.org/10.3389/fimmu.2018.00698

Liaudanskaya, V., Jgamadze, D., Berk, A. N., Bischoff, D. J., Gu, B. J., Hawks-Mayer, H., Whalen, M. J., Chen, H. I., & Kaplan, D. L. (2019). Engineering advanced neural tissue constructs to mitigate acute cerebral inflammation after brain transplantation in rats. Biomaterials, 192(November 2018), 510–522. https://doi.org/10.1016/j.biomaterials.2018.11.031

Linaro, D., Vermaercke, B., Iwata, R., Ramaswamy, A., Libé-Philippot, B., Boubakar, L., Davis, B. A., Wierda, K., Davie, K., Poovathingal, S., Penttila, P. A., Bilheu, A., De Bruyne, L., Gall, D., Conzelmann, K. K., Bonin, V., & Vanderhaeghen, P. (2019). Xenotransplanted Human Cortical Neurons Reveal Species-Specific Development and Functional Integration into Mouse Visual Circuits. Neuron, 104(5), 972–986.e6. https://doi.org/10.1016/j.neuron.2019.10.002

Luan, L., Wei, X., Zhao, Z., Siegel, J. J., Potnis, O., Tuppen, C. A., Lin, S., Kazmi, S., Fowler, R. A., Holloway, S., Dunn, A. K., Chitwood, R. A., & Xie, C. (2017). Ultraflexible nanoelectronic probes form reliable, glial scar–free neural integration. Science Advances, 3(2), 1–10. https://doi.org/10.1126/sciadv.1601966

Luhmann, H. J., Sinning, A., Yang, J. W., Reyes-Puerta, V., Stüttgen, M. C., Kirischuk, S., & Kilb, W. (2016). Spontaneous neuronal activity in developing neocortical networks: From single cells to large-scale interactions. Frontiers in Neural Circuits, 10(MAY), 1–14. https://doi.org/10.3389/fncir.2016.00040

Mansour, A. A., Gonçalves, J. T., Bloyd, C. W., Li, H., Fernandes, S., Quang, D., Johnston, S., Parylak, S. L., Jin, X., & Gage, F. H. (2018). An in vivo model of functional and vascularized human brain organoids. Nature Biotechnology, 36(5), 432–441. https://doi.org/10.1038/nbt.4127

Mcdonald, A. J. (1998). Cortical pathways to the mammalian amygdala. Progress in Neurobiology, 55(3), 257–332. https://doi.org/10.1016/S0301-0082(98)00003-3

Michelsen, K. A., Acosta-Verdugo, S., Benoit-Marand, M., Espuny-Camacho, I., Gaspard, N., Saha, B., Gaillard, A., & Vanderhaeghen, P. (2015). Area-specific reestablishment of damaged circuits in the adult cerebral cortex by cortical neurons derived from mouse embryonic stem cells. Neuron, 85(5), 982–997. https://doi.org/10.1016/j.neuron.2015.02.001

Pachitariu, M., Steinmetz, N., Kadir, S., Carandini, M., & Harris, K. (2016). Fast and accurate spike sorting of high-channel count probes with KiloSort. Advances in Neural Information Processing Systems, 1–9.

Palma-Tortosa, S., Tornero, D., Hansen, M. G., Monni, E., Hajy, M., Kartsivadze, S., Aktay, S., Tsupykov, O., Parmar, M., Deisseroth, K., Skibo, G., Lindvall, O., & Kokaia, Z. (2020). Activity in grafted human iPS cell-derived cortical neurons integrated in stroke-injured rat brain regulates motor behavior. Proceedings of the National Academy of Sciences of the United States of America, 117(16), 9094–9100. https://doi.org/10.1073/pnas.2000690117

Pham, M. T., Pollock, K. M., Rose, M. D., Cary, W. A., Stewart, H. R., Zhou, P., Nolta, J. A., & Waldau, B. (2018). Generation of human vascularized brain organoids. NeuroReport, 29(7), 588–593. https://doi.org/10.1097/WNR.0000000000001014

Qian, X., Su, Y., Adam, C. D., Deutschmann, A. U., Pather, S. R., Goldberg, E. M., Su, K., Li, S., Lu, L., Jacob, F., Nguyen, P. T. T., Huh, S., Hoke, A., Swinford-Jackson, S. E., Wen, Z., Gu, X., Pierce, R. C., Wu, H., Briand, L. A.,…Ming, G. li. (2020). Sliced Human Cortical Organoids for Modeling Distinct Cortical Layer Formation. Cell Stem Cell, 26, 1–16. https://doi.org/10.1016/j.stem.2020.02.002

Rajasethupathy, P., Sankaran, S., Marshel, J. H., Kim, C. K., Ferenczi, E., Lee, S. Y., Berndt, A., Ramakrishnan, C., Jaffe, A., Lo, M., Liston, C., & Deisseroth, K. (2015). Projections from neocortex mediate top-down control of memory retrieval. Nature, 526(7575), 653–659. https://doi.org/10.1038/nature15389

Real, R., Peter, M., Trabalza, A., Khan, S., Smith, M. A., Dopp, J., Barnes, S. J., Momoh, A., Strano, A., Volpi, E., Knott, G., Livesey, F. J., & De Paola, V. (2018). In vivo modeling of human neuron dynamics and down syndrome. Science, 1810, 1–9. https://doi.org/10.1126/science.aau1810

Rossant, C., & Harris, K. D. (2013). Hardware-accelerated interactive data visualization for neuroscience in Python. Frontiers in Neuroinformatics, 7(36), 1–9. https://doi.org/10.3389/fninf.2013.00036

Saga, Y., Miyagawa-Tomita, S., Takagi, A., Kitajima, S., Miyazaki, J. I., & Inoue, T. (1999). MesP1 is expressed in the heart precursor cells and required for the formation of a single heart tube. Development, 126(15), 3437–3447. https://doi.org/10.1242/dev.126.15.3437

Schneider, C. A., Rasband, W. S., & Eliceiri, K. W. (2012). NIH Image to ImageJ: 25 years of image analysis. Nature Methods, 9(7), 671–675. https://doi.org/10.1038/nmeth.2089

Shen, J., & Colonnese, M. T. (2016). Development of activity in the mouse visual cortex. Journal of Neuroscience, 36(48), 12259–12275. https://doi.org/10.1523/JNEUROSCI.1903-16.2016

Sweeney, M. D., Zhao, Z., Montagne, A., Nelson, A. R., & Zlokovic, B. V. (2019). Blood-brain barrier: From physiology to disease and back. Physiological Reviews, 99, 21–78. https://doi.org/10.1152/physrev.00050.2017

Tornero, D., Tsupykov, O., Granmo, M., Rodriguez, C., Grønning-Hansen, M., Thelin, J., Smozhanik, E., Laterza, C., Wattananit, S., Ge, R., Tatarishvili, J., Grealish, S., Brüstle, O., Skibo, G., Parmar, M., Schouenborg, J., Lindvall, O., & Kokaia, Z. (2017). Synaptic inputs from stroke-injured brain to grafted human stem cell-derived neurons activated by sensory stimuli. Brain, 140(3), 692–706. https://doi.org/10.1093/brain/aww347

Vivian S. Chen, James P. Morrison, Southwell, M., Foley, J. F., Bolon, B., & Elmore, S. A. (2017). Histology Atlas of the Developing Prenatal and Postnatal Mouse Central Nervous System, with Emphasis on Prenatal Days E7.5 to E18.5. Toxicology Pathology, 45(6), 705–744. https://doi.org/10.1177/0192623317728134.Histology

Wang, J., Chu, R., Ni, N., & Nan, G. (2020). The effect of Matrigel as scaffold material for neural stem cell transplantation for treating spinal cord injury. Scientific Reports, 10(2576), 1–11. https://doi.org/10.1038/s41598-020-59148-3

Wiedenmann, B., & Franke, W. W. (1985). Identification and localization of synaptophysin, an integral membrane glycoprotein of Mr 38,000 characteristic of presynaptic vesicles. Cell, 41, 1017–1028. https://doi.org/10.1016/S0092-8674(85)80082-9

Williams, R. H., & Riedemann, T. (2021). Development, diversity and death of mge-derived cortical interneurons. International Journal of Molecular Sciences, 22, 1–25. https://doi.org/10.3390/ijms22179297

Yang, G., Mahadik, B., Choi, J. Y., & Fisher, J. P. (2020). Vascularization in tissue engineering: Fundamentals and state-of-art. Progress in Biomedical Engineering, 1–18. https://doi.org/10.1088/2516-1091/ab5637

Yin, R., Noble, B. C., He, F., Zolotavin, P., Rathore, H., Jin, Y., Sevilla, N., Xie, C., & Luan, L. (2022). Chronic co-implantation of ultraflexible neural electrodes and a cranial window. Neurophotonics, 9(03), 1–13. https://doi.org/10.1117/1.nph.9.3.032204

Zakiewicz, I. M., Bjaalie, J. G., & Leergaard, T. B. (2014). Brain-wide map of efferent projections from rat barrel cortex. Frontiers in Neuroinformatics, 8(5), 1–15. https://doi.org/10.3389/fninf.2014.00005

Zudaire, E., Gambardella, L., Kurcz, C., & Vermeren, S. (2011). A computational tool for quantitative analysis of vascular networks. PLoS ONE, 6(11), 1–12. https://doi.org/10.1371/journal.pone.0027385

